# Locus-specific chromatin profiling of evolutionarily young transposable elements

**DOI:** 10.1101/2021.08.25.457666

**Authors:** Darren Taylor, Robert Lowe, Claude Philippe, Kevin C. L. Cheng, Olivia A. Grant, Nicolae Radu Zabet, Gael Cristofari, Miguel R. Branco

## Abstract

Despite a vast expansion in the availability of epigenomic data, our knowledge of the chromatin landscape at interspersed repeats remains highly limited by difficulties in mapping short-read sequencing data to these regions. In particular, little is known about the locus-specific regulation of evolutionarily young transposable elements (TEs), which have been implicated in genome stability, gene regulation and innate immunity in a variety of developmental and disease contexts. Here we propose an approach for generating locus-specific protein-DNA binding profiles at interspersed repeats, which leverages information on the spatial proximity between repetitive and non-repetitive genomic regions. We demonstrate that the combination of HiChIP and a newly developed mapping tool (PAtChER) yields accurate protein enrichment profiles at individual repetitive loci. Using this approach, we reveal previously unappreciated variation in the epigenetic profiles of young TE loci in mouse and human cells. Insights gained using our method will be invaluable for dissecting the molecular determinants of TE regulation and their impact on the genome.

## INTRODUCTION

Transposable elements (TEs) are interspersed genomic repeats that impact genome function in myriad ways (1), and whose complexity and abundance varies widely across species, as a result of intricately entangled evolutionary processes (2, 3). In humans and mice, for example, TEs make up around half of the genome, and include a wide variety of endogenous retroviruses (ERVs), long interspersed nuclear elements (LINEs) and short interspersed nuclear elements (SINEs), amongst others (4, 5). The genetic mobility of TEs, as well as their frequent involvement in recombination events, means that they are an abundant source of genetic diversity. On one hand, TE insertions can have deleterious consequences, having been implicated in a large number of genetic diseases (6) and somatic re-arrangements in cancer (7, 8). On the other hand, TE-derived sequences can be co-opted by the host genome to generate new genes (9), cis-acting regulatory elements (10), and 3D structural elements (11, 12). Moreover, transcriptionally active TEs can regulate gene in trans, functioning as long non-coding RNAs (13, 14), as well as trigger inflammatory responses through viral mimicry (15, 16).

With the growing appreciation for the impact of TEs on genomic, cellular and organismal function comes the need to have a detailed and genome-wide understanding of TE regulation. TE chromatin is tightly and dynamically regulated during development (17, 18), carefully balancing the deleterious and beneficial consequences of TE activity, with epigenetic TE deregulation being associated with developmental defects and disease (19–22). Whilst many of the generic TE chromatin regulators have been uncovered (23–25), a complete understanding of how TEs become active/silenced and impact the local genomic environment requires locus-specific information. However, the study of TE regulation has been notoriously hampered by difficulties in mapping short-read epigenomic data to individual TE copies, due to their repetitive nature (26, 27). Evolutionarily young TEs are particularly problematic, as individual copies have not diverged sufficiently from each other to allow for unique alignments across the full length of the elements. For clarity, we refer to uniquely aligned reads as those with a single best-scoring alignment, whereas non-unique reads have multiple best alignments with equal score.

One commonly used approach to retain non-unique reads from epigenomic profiling methods such as ChIP-seq, is to align reads to consensus sequences of particular TE families, thus gaining a family-wide view of epigenetic patterns. A similar outcome can be achieved by harnessing ambiguous alignments from some short-read aligners, such as bowtie2 (28), which assign non-unique reads randomly. Such ambiguously-mapped reads can be used to generate generic patterns for different TE families (see, e.g., (29, 30)). However, family-wide patterns mask differences in the epigenetic profiles of individual elements. Indeed, locus- and tissue-specific expression of young LINE-1 elements supports the notion that there is marked epigenetic variation between closely related elements of the same family (31, 32). Yet the epigenetic determinants for expression of individual TEs remain unclear. There is also a limited understanding of the locus-specific binding of transcription factors to TEs (33), which is largely limited to either evolutionarily old TEs, or to transcription factors that bind the ends of repetitive regions, benefiting from adjacent unique sequences for read alignment. Yet multiple transcription factors bind LINE-1 elements in internal regions that cannot be covered by uniquely mapped reads (29), but their contribution to locus-specific expression is unknown.

Some computational solutions have been specifically developed to assign reads from ChIP-seq data to individual repetitive elements, which use uniquely aligned reads to inform the assignment of multi-mapping reads (34, 35) (also reviewed in (26, 27)). The reliance on the existence of uniquely aligned reads places a clear limitation on the use of these methods to regions of high sequence similarity. Furthermore, whilst similar approaches are appropriate for RNA-seq data (36–38), where the signal is expected to be evenly distributed along each element, protein binding profiles are often localised and therefore difficult to detect using strategies that rely on uniquely mapped reads. Long-read solutions to mapping the epigenetic status of TEs are so far largely limited to DNA methylation profiling, which can be performed on unfragmented DNA (39). In contrast, generation of histone modification or transcription factor binding profiles traditionally require chromatin fragmentation (~300bp) in order to achieve the desired resolution.

Here we take advantage of the random 3D folding of chromatin to guide the assignment of short sequencing reads from interspersed repetitive elements (such as TEs) to individual locations. We demonstrate the accuracy of our approach using both simulated data and an orthogonal method, and showcase its ability to uncover locus-specific features of TE chromatin.

## MATERIAL AND METHODS

### Cell culture

E14 ESCs (ATCC CRL-1821) were grown in feeder-free conditions in DMEM GlutaMAX medium (Thermo Fisher) supplemented with 15% FBS, non-essential amino acids, 50 μM 2-mercaptoethanol, antibiotic-antimycotic (Thermo Fisher) and 1,000 U/ml ESGRO LIF (Millipore). MCF7 and 2102Ep cells were grown in DMEM GlutaMAX medium (Thermo Fisher), supplemented with 10% FBS, penicillin-streptomycin (Thermo Fisher) and non-essential amino acids (Gibco).

### Hi-MeDIP

Cells were crosslinked in 1% formaldehyde for 12 minutes, and the reaction quenched with 1.25 M glycine for 5 minutes. Cells were then washed three times in ice cold PBS containing protease inhibitors (Roche), flash frozen, and stored at −80°C. Thawed cells were resuspended for 30 minutes, with 5 million cells per 50 ml of ice-cold lysis buffer (10 mM Tris-HCl pH 8, 10 mM NaCl, 0.2% Igepal CA-630, one tablet protease inhibitor cocktail). After spinning down (650 *g*), cells were washed in 1x DpnII buffer, and ~5 million cells were resuspended in 716 μl of 1.32x DpnII buffer. Cells were permeabilised by adding 22 μl of 10% SDS and incubating at 37°C for 1 hour (with rotation), then quenched with 150 μl of 10% Triton X-100, followed by another 1-hour incubation at 37°C. Digestion was performed by adding 2,750 U of DpnII and incubating overnight at 37°C. After spinning down (650 *g*), the pellet was washed with 1x NEB2 buffer and resuspended in 467 μl of 1.11x NEB2 buffer. Fill-in was performed with dCTP/dGTP/dTTP (1.5 μl of each 10 mM stock), biotin-14-dATP (37.5 μl of 0.4 mM stock), and 50 U Klenow (NEB), for 1 hour at 37°C. The cell suspension was then mixed with 5 ml ligation buffer (550 μl 10x ligation buffer (NEB), 27.5 μl 20 mg/ml BSA (NEB) in water), 2,000 U of 5 U/μl T4 DNA ligase were added, and cells incubated for 4 hours at 16°C (with rotation). After DNA extraction (Allprep DNA/RNA mini kit, Qiagen), 1.5 μg DNA were sonicated to 300-700 bp using a Bioruptor Pico for 4 cycles (30 seconds on, 30 seconds off) in a total volume of 100 μL in H2O. DNA was made up to 180μL in 1x TE, and sequencing adaptors were ligated using NEBnext Ultra II reagents. For immunoprecipitation, DNA was first denatured at 95°C for 10 minutes, then cooled on ice, at which point a 20 μL input sample was taken and frozen. To the remaining DNA, 20 μL 10x IP buffer (100 mM sodium phosphate pH 7, 1.4 M NaCl, 0.5% Triton X-100) and 2 μL 5mC antibody (Active Motif 61479) were added, followed by a 2-hour incubation at 4°C. Dynabeads Protein G (previously washed in 500 μL 0.1% PBS-BSA) were resuspended in 1x IP buffer, and 8 μL beads were incubated with the DNA:antibody for 2 hours. Beads were washed twice with 350 μl 1x IP buffer for 10 minutes at room temperature, then resuspended in 125 μL Proteinase K digestion buffer (50 mM Tris-HCl pH 8.0, 10 mM EDTA, 0.5% SDS); the same was done to the stored input sample. Proteinase K (35 μg) was added and the reaction incubated at 65°C for 30 minutes. The supernatant was collected and DNA purified using SPRI beads. For purifying ligation junctions, 5 μL Dynabeads MyOne Streptavidin C1 beads (Life Technologies) were washed in 50μL Tween-buffer (5mM Tris-HCl pH 7.5, 0.5 mM EDTA, 1 M NaCl, 0.05% Tween-20) and resuspended in 83.5 μL 2x No Tween Buffer (10 mM Tris-HCl pH 7.5, 1 mM EDTA, 2 M NaCl), then mixed with 83.5 μL of DNA (150 ng maximum per 5 μL beads) and incubated at room temperature for 30 minutes. Beads were washed twice with 500 μL Tween-buffer, then once with 200 μL 1x No Tween Buffer (5 mM Tris-HCl pH 7.5, 0.5 mM EDTA, 1M NaCl), and resuspended in water. Library preparation was completed using NEBNext Ultra II reagents.

### Targeted bisulphite sequencing

ATLAS-seq (31) was adapted for bisulphite-treated DNA to comprehensively map the genomic location and assess the DNA methylation status of human full-length LINE-1 elements. The approach is focused on the youngest family (L1HS), but it also catches a significant fraction of L1PA2 to L1PA8 elements. In brief, bs-ATLAS-seq follows these steps: fragmentation of genomic DNA by sonication, ligation of a methylated adapter, bisulphite treatment of linker-ligated DNA, suppression PCR-amplification of LINE-1 5′ junctions, and asymmetric Illumina paired-end sequencing (90 bp + 210 bp). Read #1 starts from a random position (sonicated DNA break), while read #2 starts from a fixed sequence in L1 5′ UTR (internal promoter). With this design, read #2 covers the 15 most 5’ CpG sites (1-207, relative to L1HS consensus sequence). Mapping and methylation calling used bismark 0.22.1 (40).

### External datasets

The following publicly available datasets were used: mouse ESC Hi-C (GEO: GSM2533818) (41); mouse ESC H3K27ac HiChIP (GEO: GSE101498) (42); mouse AML12 Hi-C (GEO: GSE141080) and H3K9me3 HiChIP (GEO: GSE141113) (43); mouse ESC RNA-seq (GEO: GSE122854) (44); MCF7 and 2102Ep RNA-seq (EBI ENA: E-MTAB-3788) (31). More details about the sequencing datasets used in this study can be found in Supplementary Table S1.

### Hi-C and HiChIP read alignment

Data were aligned to the mm10 or hg38 genome assemblies. To perform mapping of non-unique reads informed by 3D spatial contacts, Hi-C and HiChIP data were aligned by PAtChER, with −d set between 10000 and 40000 (parameter referred to as Sd in the results section). Where not specified in the results, −d 10000 was used for mouse data, and −d 20000 for human data. After alignment, read pairing information was removed from BAM files using PAtChER’s ‘unpair’ tool. For comparison, uniquely aligned reads were extracted by filtering for ‘PO:A:u’ or ‘PO:A:s’ alignment types. To perform random mapping of non-unique reads, the ends of Hi-C and HiChIP data were independently aligned using bowtie2 (28), with default settings. Distances between interacting fragments in Hi-C data were extracted from alignments using HiCUP (45).

### RNA-seq data processing

RNA-seq data were aligned using HISAT2 (46), and reads with MAPQ lower than 2 were removed. Fragments per gene were counted with htseq-count and normalised to the total read count. Genes expression values were matched to the nearest SVA (MCF7 and 2102Ep data) or MERVL-int (mouse ESC data) element using BEDtools (47).

### Bigwig tracks

Bigwig tracks were generated using deepTools (48). Primary tracks were generated using bamCoverage, with CPM normalisation. Enrichment tracks (normalised to input) were generated using bigwigCompare, with --operation “ratio”, --pseudocount 0.01, and –skipZeroOverZero. Average profiles over selected TE families were generated with plotProfile (--averageType “mean“), and heatmap profiles with plotHeatmap (--sortUsing sum). Genome browser snapshots were generated using the WashU Epigenome Browser (49).

### Peak detection

Peaks were detected from normalised bedgraph files (converted from bigwig) using MACS2 (50) bdgbroadcall, with different values for −C depending on the experiment (see code repository for details). Peaks were intersected with either LINE-1 regions of interest for comparison with bisulphite data, or with RepeatMasker annotations to generate peak counts per TE family.

### Coverage analysis

Coverage at RepeatMasker annotations was extracted using SAMtools (51) bedcov with −d 1. Data on mouse polymorphic TEs (52) were used to remove reference B6 insertions that are absent in all sequenced 129 strains.

### Sim3C

To generate simulated Hi-C data that matched the interaction distance profile of experimental Hi-C data, a modified version of Sim3C (53) was used. In particular, the model was changed to a mix of two geometric cumulative distribution functions with different ‘shapes’ (shape1=8e-5, shape2=3e-6). The nbins parameter was set to 200,000 and alpha to 0.73 (weight of the second geometric CDF).

Minor modifications to the output format were also made. Sim3C was then run with --dist uniform, −l 75, −e DpnII, −m hic, --simple-reads, --efficiency 0.9, --anti-rate 0. The position of mapped reads was compared against that of simulated reads using custom scripts (see code repository). Files containing correctly assigned reads were used to generate accuracy tracks, which in turn were used to extract mean accuracy values per element (>1kb) of selected TE families.

### Minigenome

Sequences surrounding 960 mouse gene promoters were extracted and concatenated to make an mm10-based minigenome. A modified version of the minigenome was generated by duplicating 3kb-long regions to a position lying 3-50kb downstream from the original position (16 different distances; n=60 per distance). To assess mapping accuracy and read recovery, reads aligning to either the original or duplicate position were counted using htseq-count. Note that the duplicate regions only contain misaligned reads and no correctly aligned ones, as these regions were artificially created. Therefore, to avoid biases when normalising HiChIP data to a Hi-C control (for the enrichment and peak analyses), Hi-C reads were simulated for the duplicate regions, as follows: a) reads mapping correctly to the original location were also assigned to the duplicate location; b) reads misassigned to the duplicate location were also assigned to the original location. The same Hi-C-based reads were added to both the Hi-C and HiChIP data, to simulate non-enriched duplicate regions. Receiver operating characteristic curves for peak detection were generated by varying the −C parameter in macs2 bdgbroadcall.

### Long-range chromatin loops

To detect chromatin loops, contact matrices were generated at a 10kb resolution using Juicer (54), with hg38 reference genome and DpnII restriction sites. Significant interactions were generated using the dump command utilising the observed/expected method at a bin size of 10000. Lastly, chromatin loops were called using the HiCCUPS tool at 10kb resolution using Knight-Ruiz normalisation (54). The dump command was also used to extract the number of reads interacting within each bin involved in a chromatin loop. To assess the impact of chromatin loops on mapping accuracy, MCF7 Hi-C data were mapped to a minigenome focused on 200 chromatin loop anchors, wherein 3kb-long regions at the middle of each anchor were duplicated at the other end of the loop. For comparison, a second minigenome was made where the same regions were duplicated at a fixed distance of 50kb away.

### Hi-MeDIP data processing

To better assess the performance of PAtChER, read mappability was reduced by trimming Hi-MeDIP reads to 75bp prior to mapping. Htseq-count was used to count reads mapping to two regions of LINE-1 elements: a) a 5’ region, from 0 to 135bp from the start; b) an internal region, from 75 to 210bp from the start. Only regions containing at least three CpGs covered by the targeted bisulphite data were used. Normalised Hi-MeDIP enrichment values were compared to the mean bisulphite signal at each region.

### MERVL genomic context

MERVL-int elements were clustered based on their H3K27ac profiles, using the k-means clustering function of deepTools plotHeatmap. Distances to the nearest gene were extracted using BEDtools, and matched to their expression value. Elements from each cluster were intersected with a published annotation of spatial A and B compartments (41).

## RESULTS

### 3D chromatin information improves mapping coverage at TEs

The use of 3D conformation data from Hi-C experiments has been improving the assembly of genomes for nearly a decade now (54). We argued that a similar principle could be employed to enable the assignment of non-unique short sequencing reads to specific locations, provided the source material was genomic DNA. In particular, we hypothesized that coupling of ChIP-seq to Hi-C (55, 56) would enable a chromatin conformation-guided mapping of each end of the sequencing library to produce a ChIP-seq track with increased coverage of interspersed repeats. As Hi-C generates chimeric fragments between genomic regions lying nearby in 3D space, non-unique reads can become paired with unique reads from proximal genomic regions (Figure 1A). A non-unique end can therefore be assigned with a relatively high degree of confidence to the mapping location that lies nearest to the corresponding unique end (Figure 1A), given the steep inverse proportional relationship between genomic distance and contact frequency (Figure 1B). Importantly, high frequency contacts within a few tens of kb are largely due to random collisions, whereas non-random interactions can normally only be discerned from the latter at distances over 50 kb (57–59).

**Figure 1.**
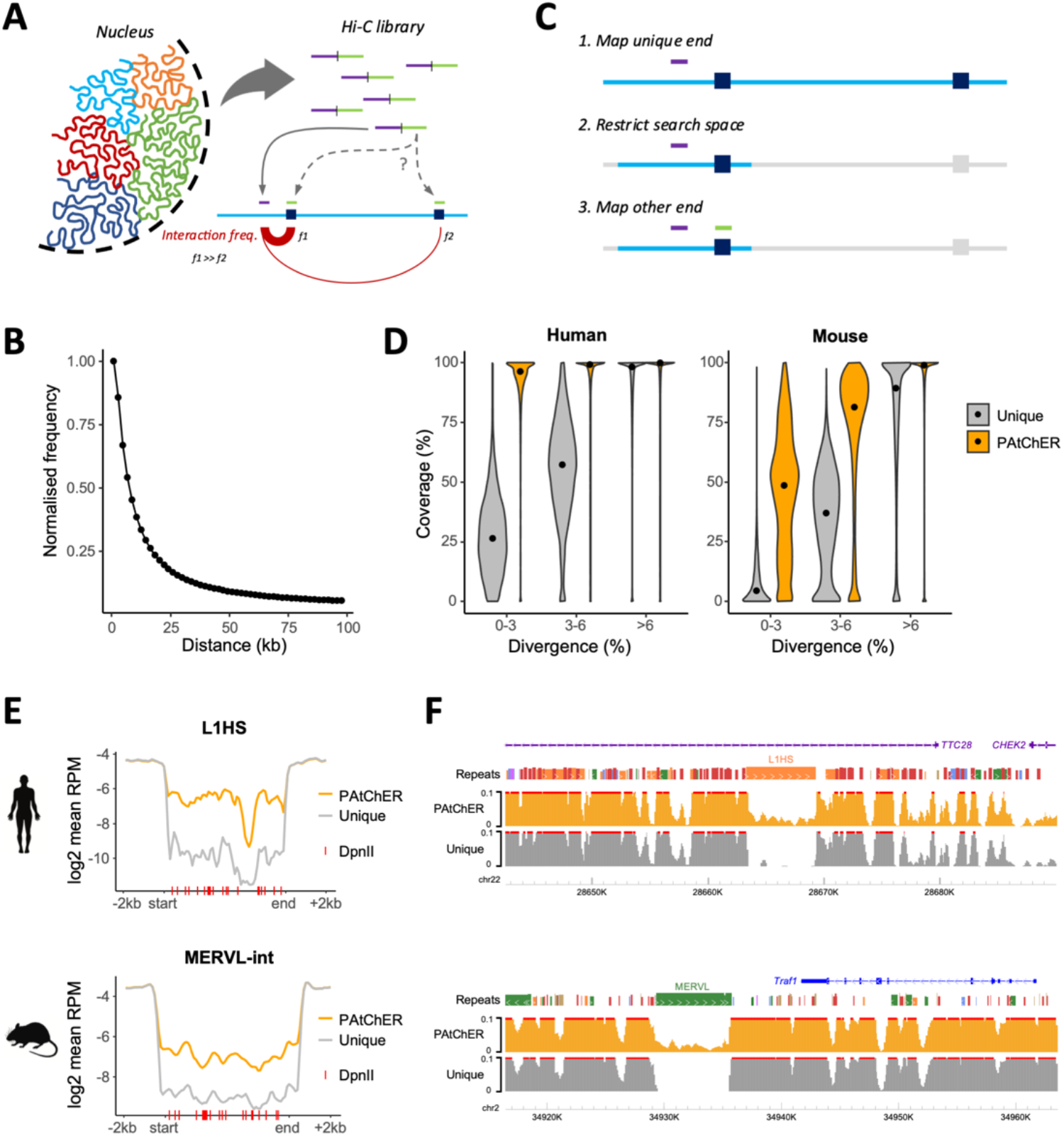
Harnessing Hi-C data to increase sequencing coverage at interspersed repeats. (**A**) Hi-C libraries generate chimeric reads between spatially close genomic fragments. Uniquely mapping DNA fragments (purple) are more likely to interact with repetitive fragments (green) from nearby loci, due to the high frequency of random collisions between tethered loci. (**B**) Frequency of 3D interactions as a function of inter-locus genomic distance, derived from mouse ESC Hi-C data. (**C**) The mapping principle of PAtChER. Hi-C fragments with one uniquely mappable end (purple) and one multimapping end (green) are processed by first aligning the unique end; a hit for the non-unique end is then sought within a user-defined distance (Sd) from the unique end. (**D**) Coverage (% of bp covered by at least 1 read) of repetitive elements larger than 1kb as a function of their sequence divergence to the respective consensus sequence. Hi-C data from human or mouse cells was processed by PAtChER, and compared against uniquely mapped reads alone. (**E**) Average sequencing depth across L1HS (>5kb) or MERVL-int (>4kb) elements in PAtChER- or uniquely-aligned data. (**F**) Genome browser snapshots showing examples of sequencing dept at L1HS (top) and MERVL (bottom) elements.

To leverage this alignment principle, we developed PAtChER (<u>P</u>roximity-based <u>A</u>lignment of Hi<u>Ch</u>IP <u>E</u>nds to <u>R</u>epeats; https://github.com/MBrancoLab/PAtChER), a computational pipeline to map the ends of HiChIP data to produce ChIP-seq tracks with increased genomic coverage. When PAtChER encounters Hi-C fragments for which only one end can be uniquely aligned, it uses its location as an ‘anchor’ to search for possible mapping hits of the non-unique end in its vicinity (Figure 1C). Instead of searching for all possible hits of the non-unique end in the genome (which would use up prohibitive amounts of processing time), PAtChER restricts the search space by only looking for hits within a user-defined distance from the unique end (Figure 1C). If more than one hit is found within this search space, the nearest hit to the unique end is chosen. The search distance parameter (which we will denote here as Sd) therefore affects the stringency of the mapping, as hits beyond this distance will not be considered.

We evaluated the gains in mapping coverage by using PAtChER (Sd=20kb) to process human and mouse Hi-C data – our own Hi-C dataset from 2102Ep cells, and a published dataset from mouse ESCs (41). We focused on repeats longer than 1kb, and segregated them based on the degree of divergence to the respective family consensus (which also functions as a proxy to their evolutionary age). The fraction of each element that is covered by at least 1 read was measured and, as expected, lowly divergent repeats were poorly covered by uniquely mappable reads (Figure 1D).

Including reads recovered by the PAtChER mapping strategy led to a dramatic increase in the coverage of these young repeats, especially in human data (Figure 1D). The lower mappability of young mouse repeats relative to human ones is likely due to their higher abundance, as previously suggested (60), which reduces the number of unique ‘anchor’ points to support PAtChER-based mapping. When focusing on specific TE families, we found that PAtChER-aligned data virtually fully covered young LINE-1 and SVA elements in human (Supplementary Figure S1A). Coverage in mouse TEs displayed a bimodal distribution at some young TE families (Supplementary Figure S1B). When we excluded polymorphic elements known to be absent in the 129 strain (52) (from which E14 ESCs were derived), the distribution became largely unimodal, with IAPEz elements displaying a median coverage of 74% (Supplementary Figure S1C). Notably, unlike for uniquely mapped reads, coverage by PAtChER-mapped reads increases substantially with sequencing depth, due to an increased probability of capturing informative 3D contacts (Supplementary Figure S1D).

Coverage by PAtChER-mapped data was generally even across TEs, although this can be affected by the density of restriction sites used during Hi-C library preparation (Figure 1E, note dip in L1HS coverage at a DpnII-poor region). Importantly, read density was markedly higher than with uniquely mapped data alone, even though it was not as high as in adjacent non-TE regions, as expected (Figure 1E,F). To deal with this uneven read depth, when applying PAtChER to HiChIP data, it is important to include a Hi-C input control in order to normalise the signal (Figure 3A; see below). This allows quantification of enrichment for proteins of interest at regions that would otherwise be excluded from sequencing data.

### PAtChER maps non-unique reads with high accuracy

To test whether PAtChER assigns reads to their correct locations, we first simulated Hi-C data using Sim3C (53). Importantly, we adjusted the simulated model to ensure that the interaction distance distribution matched that of our own Hi-C data (Supplementary Figure S2A). We used PAtChER to map human and mouse Sim3C-generated data using different values of Sd. To determine accuracy, we calculated the percentage of PAtChER-mapped reads that were assigned to the correct (simulated) location. Overall, PAtChER yielded high mapping accuracy of non-unique reads (>75%), with better performance in human than in mouse data (Figure 2A). Whilst increasing the mapping search distance (Sd) decreased PAtChER accuracy, it led to gains in read recovery (Figure 2A). We therefore sought to evaluate the trade-off between accuracy and read recovery at specific TE families. All human TE families tested and several mouse ones (e.g., MERVL, IAPEz, ETnERV) benefited from the increased read recovery that come with higher values of Sd, without compromising substantially on accuracy (Figure 2B; Supplementary Table S2). In contrast, accuracy at young mouse LINE-1 families dropped rapidly with higher Sd values, arguing for the need to carefully choose Sd based on the target TE families of interest. Elements lying in repeat-dense regions, such as IAPEy-int (clustered on the Y chromosome) and certain Alu-dense clusters, have notably reduced read recovery likely due to a lower number of unique anchor points, yet maintain high mapping accuracy (Supplementary Table S2). Mapping accuracy also varied between copies of the same family (Supplementary Figure S2B), depending on the local genomic environment. To aid the quality control of results we have generated genome-wide mapping accuracy tracks (Figure 2C; Supplementary Figure S2C; Supplementary Table S3). This also enables an appreciation of how accuracy changes within an element, as seen for LINE-1 elements, wherein the 5’ end maps more accurately than the 3’ end, as expected from the abundance of truncated ORF2 fragments within the human genome (Figure 2D; Supplementary Figure S2D).

**Figure 2.**
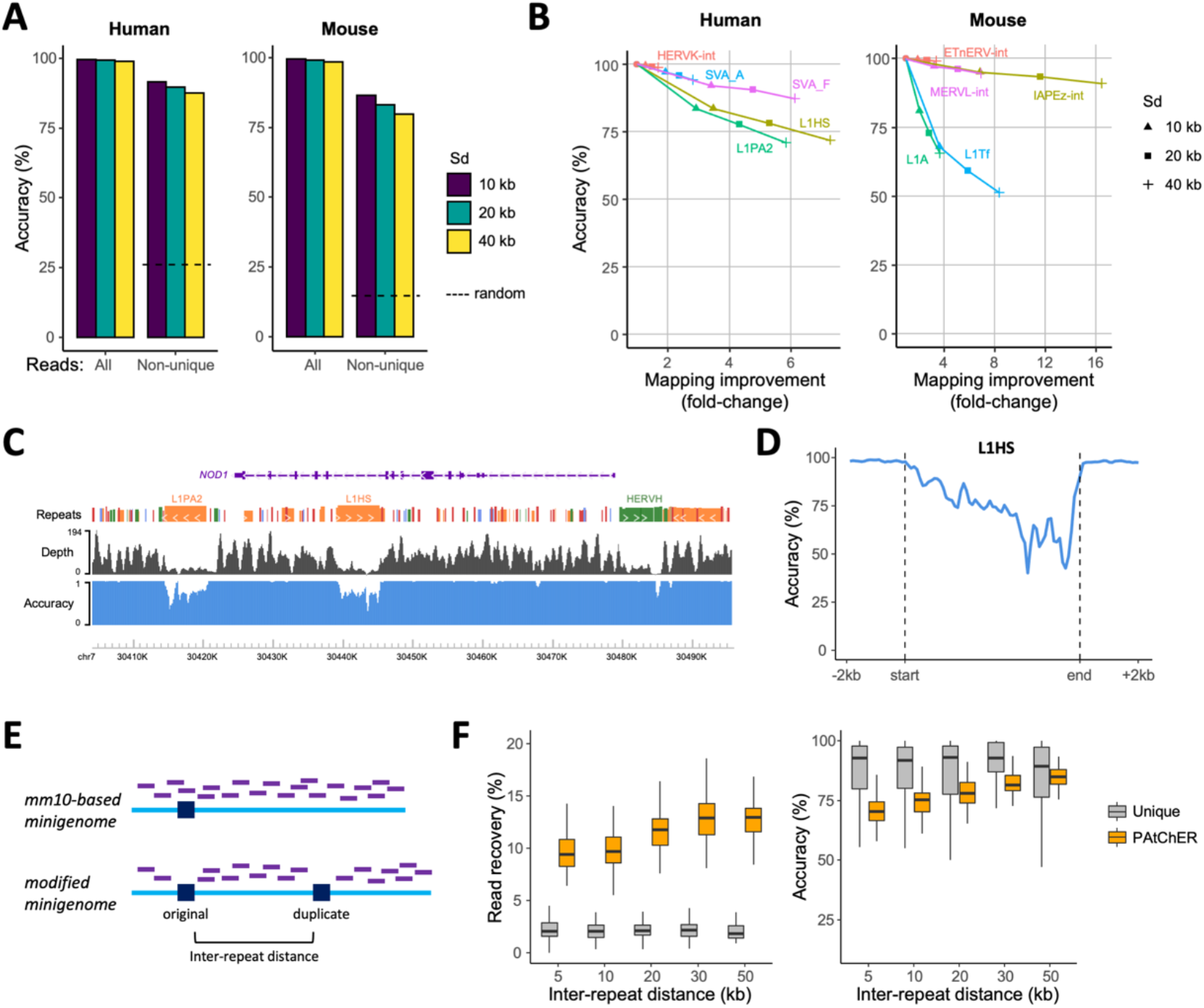
PAtChER mapping accuracy. (**A**) Overall mapping accuracy of Sim3C-simulated reads. Human and mouse Hi-C data were simulated and mapped with PAtChER using different values of Sd. For comparison, the accuracy for randomly assigned non-unique reads (by bowtie2) is shown (dashed line). (**B**) The change in mapping accuracy of Sim3C data was compared against the change in number of mapped reads at selected TE families (elements >1kb) and for different Sd values. Read numbers are normalised against the value obtained using unique alignments (circles). (**C**) Genome browser snapshot of read depth and mapping accuracy tracks for the human genome with Sd=20kb, using Sim3C data. (**D**) Average mapping accuracy profile across L1HS (>5kb) elements, using Sim3C data. (**E**) Principle of the minigenome-based approach to assess mapping accuracy. Hi-C data was mapped to a minigenome made up of 960 merged genomic segments, which contained either mm10 reference sequences, or were modified to include 3kb-long duplications at a variable ‘inter-repeat’ distance from their original locations. Reads (purple lines) that cover the duplicated regions in the mm10-based minigenome become unmappable uniquely in the modified genome. (**F**) Hi-C data mapped to the minigenomes (Sd=20kb) were assessed in terms of read recovery (% of reads mapped at original location compared to a mm10-based minigenome alignment) and mapping accuracy (% of reads mapped to original, correct location), as a function of inter-repeat distance.

We also evaluated mapping accuracy using real Hi-C data, by aligning reads to a ‘minigenome’ containing artificially duplicated 3kb-long regions, spaced out by variable inter-repeat distances (Figure 2E). By comparing with alignments to a non-modified, mm10-based version of the minigenome (i.e., without the duplications), we could quantify how many reads were recovered, and how many were assigned to the original (correct) or duplicate (incorrect) locations. Read recovery (% of reads at original location in the modified minigenome over those in the mm10-based minigenome) was dramatically higher in PAtChER-mapped data when compared with unique reads, and increased marginally with higher inter-repeat distance (Figure 2F; Sd=20kb). Mapping accuracy was in the same high range as seen in Sim3C data, and expectedly increased with increasing inter-repeat distance, reaching levels comparable to uniquely mapped reads when the gap was 50kb (Figure 2F; Sd=20kb). Other values of Sd provided trade-offs between accuracy and read recovery (Supplementary Figure S2E), as already highlighted above.

Finally, we asked whether mapping accuracy could be affected by non-random long-range interactions, which play important roles in the spatial and functional organisation of genomes (62). Long-range chromatin loops involving two TEs of the same family could lead to read misassignment. We therefore performed chromatin loop detection in two human Hi-C datasets (MCF7 and 2102Ep cells), and compared the number of reads involved in chromatin loops with those involved in nearby random contacts. Irrespective of the size of the interacting region, proximal contacts outweighed the number of reads involved in chromatin loops by around 12-fold (Supplementary Figure S3A,B), suggesting that chromatin loops are unlikely to significantly affect mapping accuracy. To confirm this, we generated a new minigenome wherein duplicated regions were placed at the base of chromatin loops (Supplementary Figure S3C). Reassuringly, mapping accuracy remained high and similar to that seen in a minigenome with duplicated regions placed 50kb apart (Supplementary Figure S3D).

These results provided us with confidence that PAtChER maps reads accurately, provided that adequate values of Sd are chosen in a species- and TE family-specific manner.

### Accurate protein enrichment detection at repetitive regions using Hi-ChIP and PAtChER

Our main objective was to use PAtChER to map Hi-ChIP data in a manner that produces ChIP-seq-like profiles at interspersed repeats. To achieve this, it is critical to include a control Hi-C library (equivalent to the input in ChIP-seq experiments) that is used to control for uneven sequencing depth (Figure 3A). To test the quality and accuracy of data processed in this manner, we again mapped data to the minigenome containing artificially duplicated regions. We used H3K27ac Hi-ChIP data from mouse ESCs (42), and normalised it to Hi-C data from the same cell line (41). Visual inspection of normalised data showed that PAtChER was able to reconstruct regions of H3K27ac enrichment at the original location, whereas the duplicate region had a much reduced signal (Figure 3B). In contrast, no enrichment was seen at either region using only uniquely mapped reads (Figure 3B).

**Figure 3.**
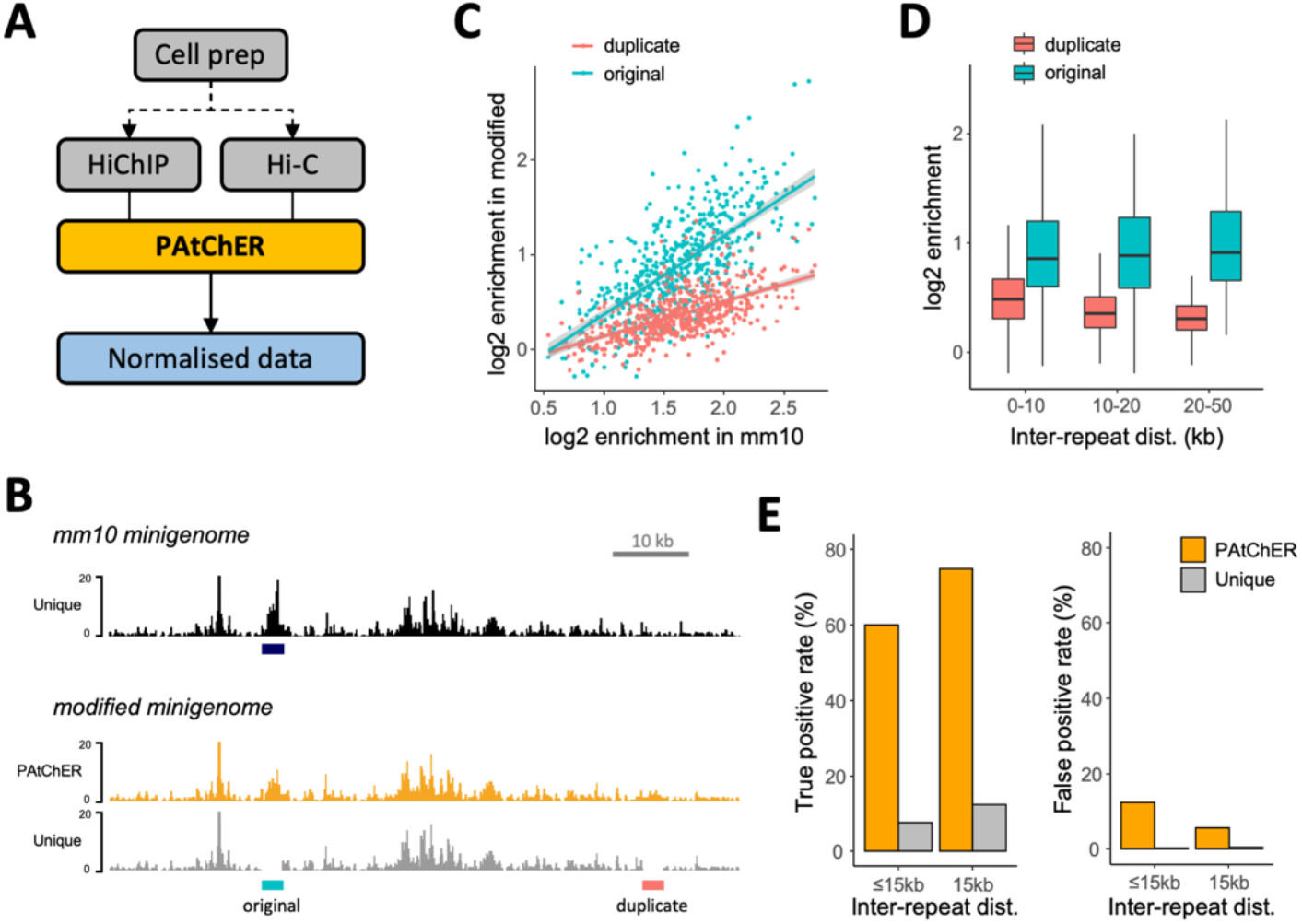
Accuracy of protein enrichment detection at repeats. (**A**) To produce ChIP-seq-like profiles from HiChIP data, parallel processing by PAtChER of a Hi-C input is essential to allow for normalisation of uneven coverage. This can be done from the same cell prep (after cross-linking, digestion, ligation) by dividing it into two before the immunoprecipitation step. (**B**) H3K27ac HiChIP data were mapped to the mm10-based and modified minigenomes (see Figure 2E), and normalised to Hi-C input data. Genome browser snapshot demonstrates how PAtChER recreates a peak of H3K27ac enrichment at the original position, as seen in the mm10 minigenome. (**C**) H3K27ac enrichment values at original (correct) and duplicate (incorrect) locations in the modified minigenome, and compared with the enrichment at the original location in the mm10 minigenome. (**D**) H3K27ac enrichment values at original and duplicate locations as a function of inter-repeat distance. (**E**) Peak detection performance at modified minigenome as a function of inter-repeat distance. The true positive rate is the % of mm10 peaks that are found at original location in the modified minigenome. The false positive rate is the % of duplicate locations where peaks were detected.

We extended this analysis to all 960 duplicated regions within the minigenome, first by calculating enrichment values at the both original and duplicate locations, and comparing them to enrichment values from mapping to the non-modified, mm10-based minigenome. Enrichment values at the original location of the modified minigenome correlated well with those from the mm10-based minigenome (Figure 3C), showing that normalisation of HiChIP PAtChER-aligned data enables detection of enriched regions at interspersed repeats. Importantly, enrichment values were distinctly lower at the duplicate (incorrect) locations, with the difference increasing for higher enrichment values (Figure 3C). This difference also increased with higher inter-repeat distance (Figure 3D), as expected from the higher mapping accuracy when repeats are further apart. To test how discriminatory these differences in enrichment were, we performed peak detection on normalised data using MACS2 (50). For optimised peak detection conditions, we found that PAtChER enabled recovery of 60-75% of the expected peaks, depending on inter-repeat distance (Figure 3E). The false discovery rate for repeats separated by more than 15kb was 5.6%, showing that peak detection is highly accurate for this group (Figure 3E). By varying a key peak detection parameter, we calculated an overall area under the curve (AUC) of 0.95 for repeats >15kb apart, whereas uniquely mapped data had an AUC of only 0.73 (Supplementary Figure S4).

Overall, these analyses suggest that PAtChER processing of HiChIP data enables accurate detection of genomic regions of protein enrichment.

### Validation of PAtChER outputs using an orthogonal method

As a final validation of PAtChER-aligned data at TEs, we compared two orthogonal methods to map DNA methylation at interspersed repeats: 1) a combination of Hi-C with methylated DNA immunoprecipitation (i.e., Hi-MeDIP), followed by processing by PAtChER ; and 2) bisulphite sequencing of the promoter of young human LINE-1 elements, using a modified version of ATLAS-seq (31) that provides locus-specific DNA methylation levels of the most distal 210 bp of full-length LINE-1 elements. We performed Hi-MeDIP and LINE-1 bisulphite sequencing on MCF7 (breast cancer) and 2102Ep (embryonal carcinoma) cell lines. After PAtCHER alignment and normalisation of the Hi-MeDIP data, we used only non-unique reads to measure the enrichment at the most internal part of the LINE-1 region covered by bisulphite sequencing (Figure 4A). As a comparison, we also analysed the more mappable 5’ end of the bisulphite-covered region, calculating Hi-MeDIP enrichments therein using only unique reads (Figure 4A). We found a good correlation between Hi-MeDIP data and bisulphite sequencing irrespective of whether unique (5’ end) or non-unique (internal region) reads were used, showing good accuracy of PAtChER-processed enrichment values (Figure 4B). We obtained a similar outcome when including both unique and non-unique reads (Supplementary Figure S5A).

**Figure 4.**
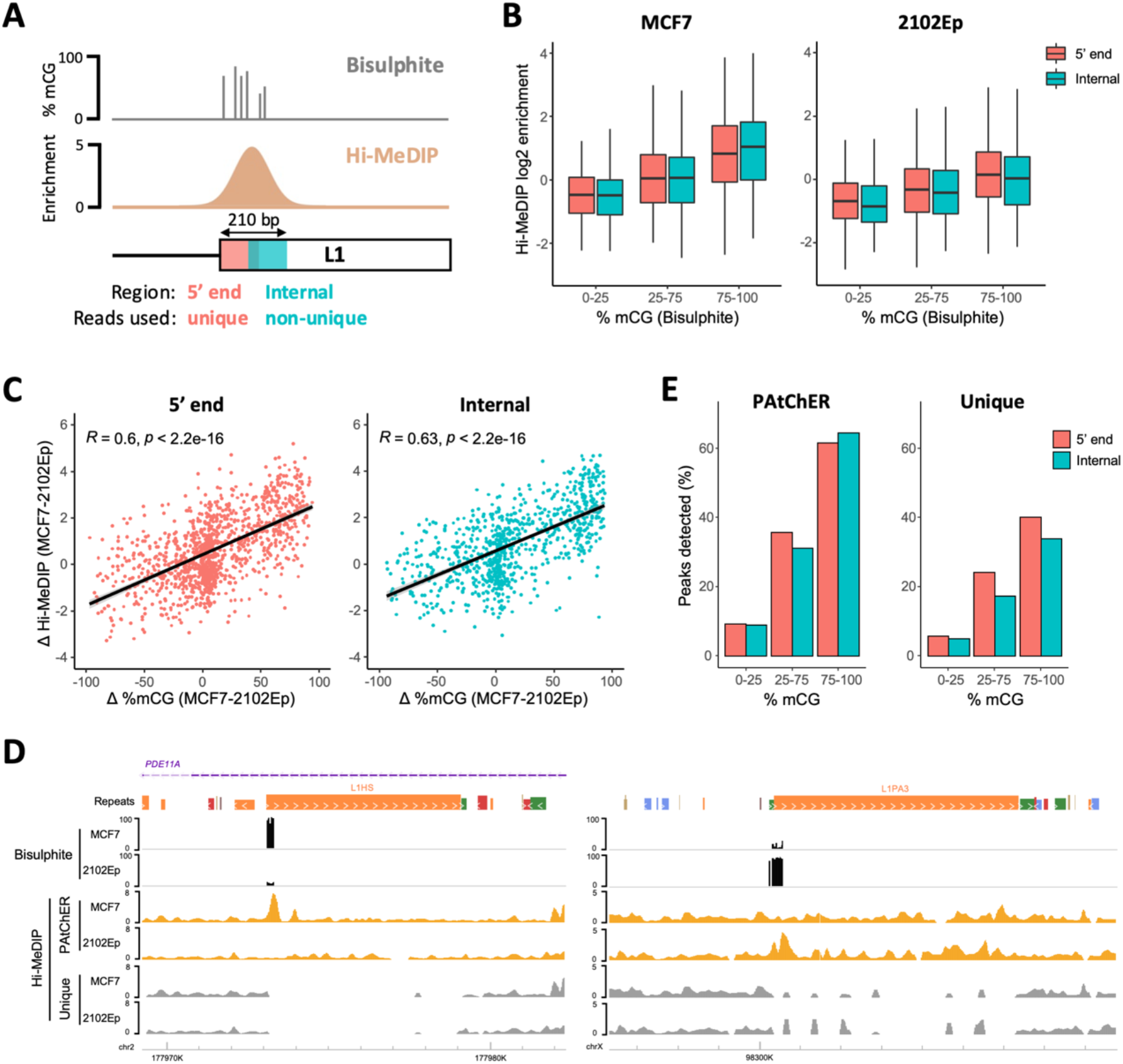
Validation of PAtChER-processed data using an orthogonal method. (**A**) DNA methylation of human LINE-1 elements was profiled using Hi-C combined with antibody-based enrichment of DNA methylation (Hi-MeDIP) and processing by PAtChER, and compared to data from a targeted bisulphite sequencing approach that measures DNA methylation at the first 210 bp of LINE-1 elements. Enrichment values were calculated using either unique reads from the 5’ end, or using non-unique reads from a less mappable internal region. (**B**) Correlation between PAtChER-processed Hi-MeDIP enrichment values and bisulphite sequencing data (expressed as % of methylated CpGs) at both LINE-1 regions, in MCF7 and 2102Ep cells. (**C**) The DNA methylation difference between MCF7 and 2102Ep cells was calculated using data from either Hi-MeDIP or bisulphite sequencing and compared, yielding a strong correlation for both LINE-1 regions (Pearson’s R is displayed). (**D**) Peak detection was performed on PAtChER-processed or uniquely aligned data. The plots display the percentage of peaks overlapping each LINE-1 element region, depending on their bisulphite methylation status. (**E**) Genome browser snapshots displaying bisulphite and Hi-MeDIP data at LINE-1 elements with preferential methylation in either MCF7 (left) or 2102Ep (right) cells.

It is well established that the signal in MeDIP experiments is affected by sequence composition, adding noise to comparisons between different genomic regions. A more quantitative approach is to analyse the difference in signal between two samples at the same loci. We therefore calculated the difference in enrichment between MCF7 and 2102Ep, which yielded a strong correlation with bisulphite data (Figure 4C; Supplementary Figure S5B). Importantly, the correlation between the two techniques is very similar when comparing uniquely aligned data (at the 5’ end) with PAtChER-aligned non-unique reads (at the internal region; Figure 4C). Examples of individual loci showcase the robustness in PAtChER-derived enrichment values over uniquely mapped data (Figure 4D).

Finally, we performed peak detection on the normalised data, considering either all PAtChER-aligned reads, or only unique ones. In both cases, peaks were rarely detected in regions of low bisulphite signal, with most peaks lying in hypermethylated regions with >75% methylation (Figure 4E). Unique reads were sufficient to detect peaks at 40% of hypermethylated regions at the 5’ end, where mappability is relatively high, but this value was lower within the less mappable internal region of LINE-1s (Figure 4E). Using PAtChER-aligned data led to a clear improvement in peak detection over uniquely aligned data, with 62-64% of hypermethylated regions overlapping with peaks (Figure 4E). Notably, PAtChER-aligned data performed equally well in both LINE-1 regions, showing that internally detected peaks are of high confidence.

This genome-wide orthogonal validation using a highly quantitative technique demonstrated that PAtChER data provides reliable detection and quantification of protein enrichment at evolutionarily young interspersed repeats.

### PAtChER-processed data reveals TE copy-specific epigenomic profiles

Having validated PAtChER-processed data through multiple approaches, we sought to gain insights into TE chromatin profiles at the level of individual elements. Our Hi-MeDIP analysis had already revealed locus- and tissue-specific patterns of LINE-1 DNA methylation that could not be appreciated from uniquely mapped reads alone (Figure 4E). To expand this analysis, we looked for repeat families wherein more DNA methylation peaks could be detected when comparing PAtChER-processed reads with uniquely aligned reads. Apart from young LINE-1 elements, a number of other young TE families were associated with DNA methylation peaks, including SVA_F elements (Figure 5A). Indeed, most SVA elements larger than 1kb displayed localised DNA methylation enrichment that could not be appreciated from uniquely mapped data (Figure 5B,C). SVA elements are known to harbour regulatory activity, including in pluripotent cells (63), suggesting that their hypomethylated status in 2102Ep embryonal carcinoma cells (when compared to MCF7) could be related to gene regulatory activity. However, we found no evidence of an overall association with the transcriptional activity of neighbouring genes (Supplementary Figure S6A). It may be that specific elements do play regulatory roles, such as in the well-characterised example of the PARK7 locus (64) (Supplementary Figure S5B), but testing this possibility would require genetic and/or epigenetic editing experiments (63). Having copy-specific chromatin profiles can thus enable the identification of candidate gene regulatory loci. We also generated Hi-MeDIP data for mouse ESCs, which revealed a larger number of DNA methylation peaks at young TE families when comparing PAtChER processing to uniquely mapped data, including at IAPEz and RLTR10 elements (Supplementary Figure S6C,D).

**Figure 5.**
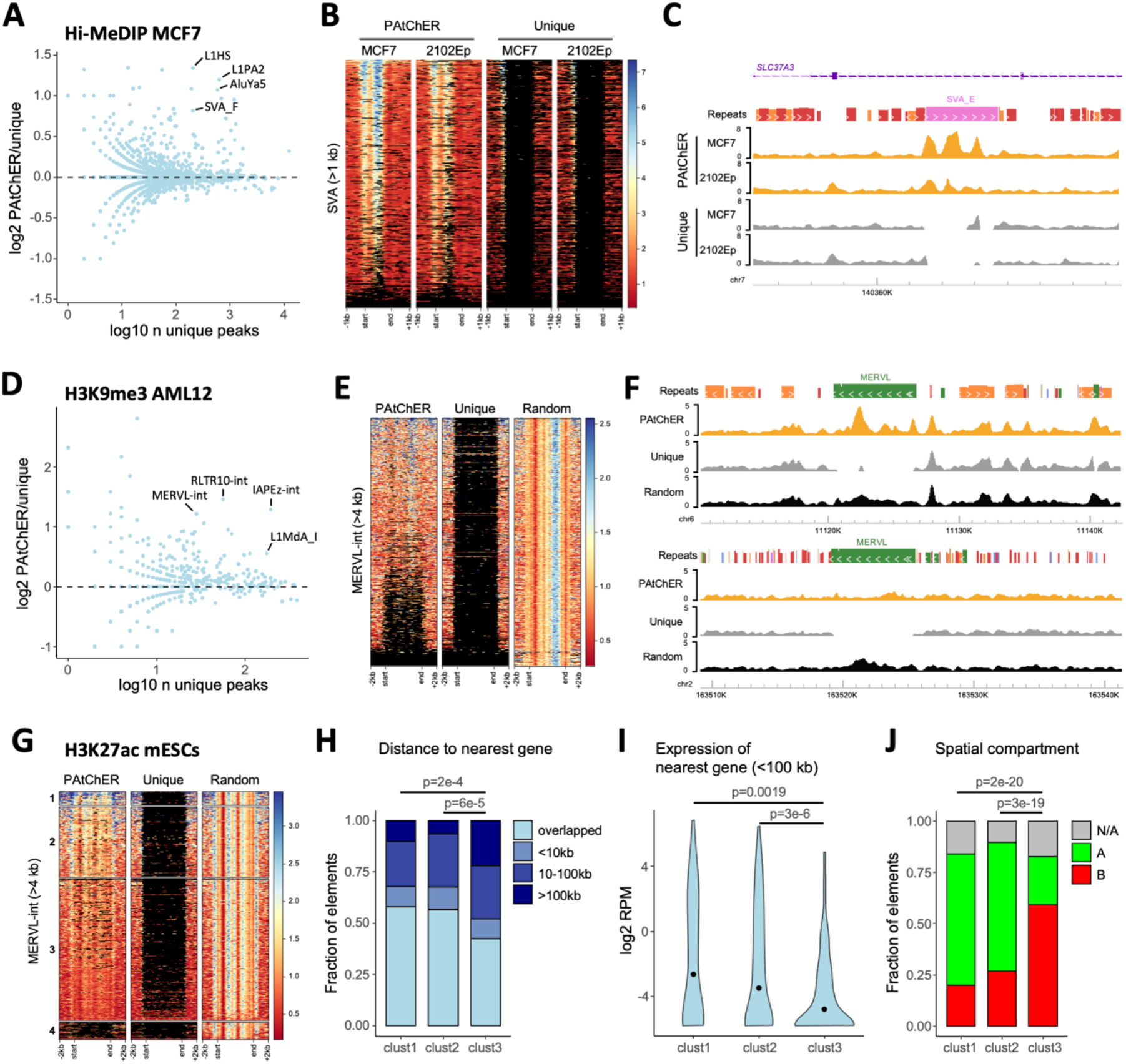
Revealing TE chromatin variation using HiChIP and PAtChER. (**A**) The number of DNA methylation peaks (from Hi-MeDIP) detected in MCF7 for each TE family was compared between PAtChER- and uniquely-aligned data. (**B**) DNA methylation enrichment profiles at SVA elements larger than 1kb in MCF7 and 2102Ep cells. Each line in the heatmap represents a single SVA locus. (**C**) Genome browser snapshot of an example SVA locus where the DNA methylation profile can only be appreciated from PAtChER-aligned data. (**D**) Comparison of PAtChER and unique H3K9me3 peaks overlapping different TE families in mouse AML12 cells. (**E**) H3K9me3 enrichment profiles at MERVL-int elements larger than 4kb in AML12 cells. PAtChER and uniquely aligned data are compared with bowtie2-aligned data, where non-unique reads are randomly assigned a best location. (**F**) Genome browser snapshots of MERVL loci where clear differences in H3K9me3 enrichment are visible that cannot be appreciated from uniquely- or bowtie2-aligned data. (**G**) H3K27ac enrichment profiles at MERVL-int elements larger than 4kb in mouse ESCs. K-means clustering divided the elements into highly (1) or moderately (2) enriched, unenriched (3), and no covered (4). (**H**) Distance of MERVL elements in each cluster to annotated genes. P-values are from chi-squared tests, corrected for multiple comparisons using the Benjamini-Hochberg method. (**I**) Expression distribution of genes within 100kb of MERVL elements in each cluster. P-values are from Wilcoxon tests, corrected for multiple comparisons. (**J**) Overlap of MERVL elements in each cluster with annotated A/B spatial compartments in mouse ESCs. N/A indicates elements overlapping unannotated regions. P-values are from chi-squared tests, corrected for multiple comparisons.

We then processed published HiChIP data for H3K9me3 (43), a major TE-silencing mark that often displays a broad distribution that is difficult to assess from uniquely mapped reads (65). Similar to DNA methylation in ESCs, H3K9me3 in AML12 hepatocytes showed a higher number of peaks at young TE families when processed by PAtChER (Figure 5D). Many of these peaks were located deep within the respective element, in areas that are not covered by uniquely mapped reads, as exemplified by the distribution of H3K9me3 at MERVL elements (Figure 5E,F). Mapping approaches that assess TE family-wide patterns can reveal such internally located peaks, but fail to capture the variability between individual elements. Indeed, when we mapped the data using bowtie2 (28), which randomly assigns non-unique reads, enrichment hotspots were found evenly distributed across MERVL copies (Figure 5E,F). In contrast, PAtChER captured differences in H3K9me3 deposition between elements of the same family, revealing epigenetic differences that may affect or be affected by the local environment.

Finally, we investigated the distribution of an active histone mark (H3K27ac) in mouse ESCs, where several TE families are known to be active and play important roles (14, 44, 66). Using deep H3K27ac HiChIP data (42), we compared PAtChER-aligned reads with unique alignments, and random assignment of non-unique reads. Of particular notice was the MERVL family, which after PAtChER processing displayed marked variation in H3K27ac enrichment between individual elements, including within the internal coding regions, which could not be appreciated by the other two approaches (Figure 5G). MERVL elements are highly active during the 2-cell stage of mouse preimplantation development, and are thought to facilitate zygotic genome activation via gene regulatory activity (67–69). Although MERVL elements are only transcribed in a small proportion of ESCs (68), our data showed that a subset of elements are enriched in H3K27ac (Figure 5G). We therefore asked whether H3K27ac-enriched elements were embedded within a distinct genomic environment. K-means clustering divided elements into groups with high enrichment (cluster 1), moderate enrichment (cluster 2), and low to no enrichment (cluster 3) (Figure 5G). A fourth cluster contained few aligned reads, and may constitute polymorphic elements that are not present in the 129 genome of E14 ESCs. A slightly higher proportion of elements from enriched clusters (cluster 1 and 2) we positioned intragenically when compared with cluster 3 elements (Figure 5H). A more prominent difference was found when we analysed the expression levels of genes within 100kb of MERVL elements, with those lying near cluster 1 and 2 being substantially more active than genes near cluster 3 elements (Figure 5I). Accordingly, cluster 1 and 2 MERVL elements were predominantly found within the so-called A compartment of spatial genome organisation (characterised by euchromatic, active regions), whereas most cluster 3 elements were found within the B compartment (characterised by heterochromatic, inactive regions) (Figure 5J). This suggests that in ESCs, the surrounding chromatin and 3D environments are major determinants of MERVL epigenetic status. Therefore, the dramatically distinct landscape of the 2-cell stage (70–73) may facilitate MERVL expression and enable its functional roles therein.

## DISCUSSION

We have presented here a strategy for generating protein-DNA binding profiles at interspersed repeats using data from HiChIP or similar methods. HiChIP is now a well-established technique that is used by an increasing number of laboratories, and for which commercial kits are available. Therefore, the experimental implementation of our strategy is already well supported, and the only additional requirement is the use of PAtChER to process the data, making our solution highly accessible to other researchers.

We have shown through multiple approaches that PAtChER-processed data is accurately mapped. Whilst the level of accuracy varies between species, TE family, and even individual elements, using different values of Sd and filtering data based on mapping accuracy information can help to ensure that results are reliable. It is also worth noting that, when detecting areas of enrichment, any mis-aligned reads are more likely to decrease sensitivity than yield false positives because, unlike areas of true enrichment, alignment errors will be randomly distributed. This prediction is supported by the low rate of Hi-MeDIP peak detection at hypomethylated human LINE-1 elements (Figure 4D). Peak detection using the minigenome approach also displays a low false positive rate (Figure 3E), and we believe this to be a conservative estimate, because here the random spread of mis-aligned reads is limited by the fact that there are only two copies of each repeat. We propose that the use of similar minigenomes (i.e., pseudo reference genomes) could be used in a flexible and bespoke manner to estimate false positive rates from any HiChIP dataset, taking into account the size and spacing of TEs of interest. We have also shown that long-range chromatin loops have no discernible effect on mapping accuracy, due to their much lower contact frequency when compared to proximal random interactions (Supplementary Figure S3). Nonetheless, the potential bias generated by cases of particularly strong interactions between repeats should not be overlooked. These specific cases can be assessed by extracting and analysing long-range interactions from the Hi-C or HiChIP data used for PAtChER-based mapping.

A number of changes can be employed to the workflow presented. Namely, all the HiChIP datasets analysed here relied on restriction enzymes for the initial chromatin digestion, which can lead to uneven coverage, as seen for L1HS elements (Figure 1E). This limitation can be easily overcome by the use of existing restriction-independent digestion strategies (74, 75), which result in more even coverage of the genome. We also note that to attain adequate coverage of young TEs requires a relatively large amount of sequencing for large genomes (Supplementary Figure S1D).

These sequencing requirements are similar to those of a high-resolution HiChIP experiments where the objective is to map 3D interactions. Nonetheless, we suggest that when the objective is to apply PAtChER solely to obtain ChIP-seq-like profiles, sequencing requirements can be reduced by the use of capture probes to specific TE families (76), thus focusing the sequencing efforts on the regions of interest. Finally, similar to essentially all past ChIP-seq experiments, we have restricted our analyses to elements annotated within reference genomes. A more accurate assessment of the TE epigenomic landscape would require taking into account variation at polymorphic TE loci. Interestingly, our coverage analysis showed that reads are not mis-assigned to reference B6 TEs that are absent in the 129 mouse genome (Supplementary Figure S1C), suggesting that PAtChER can identify these ‘absent’ variants. Moreover, spatial genomic contacts can also be used to identify non-reference TE insertions, as previously shown (76). It may therefore be possible to design computational strategies to analyse HiChIP data in a manner that delivers epigenomic profiles at non-reference insertions, although not having complete sequences for those insertions constitutes a significant limitation.

Recently, a solution for profiling protein-DNA interactions based on long-read Oxford Nanopore Technologies (ONT) sequencing was proposed, termed DiMeLo-seq (77). It combines antibody-directed local DNA adenine methylation, followed by ONT sequencing. Whilst TEs were not analysed in this study, and therefore the robustness and validity of the data at interspersed repeats remain unclear, DiMeLo-seq nonetheless promises in principle to be a viable long-read-based alternative to our short-read-based approach. The two different approaches have a different set of advantages and disadvantages. We have listed some of the current limitations of our approach above. Regarding DiMeLO-seq, it critically depends on efficient in situ DNA methylation, and requires whole genome ONT sequencing, which is currently expensive. Additionally, ONT sequencing remains relatively error prone when compared to short-read sequencing, and requires large amounts of input DNA – unlike HiChIP, which can be performed on small cell numbers (42). Therefore, we believe that the existence of complementary approaches is important, and will enable cross-validation of results.

Using PAtChER-processed HiChIP data, we have demonstrated that TEs harbour a large degree of epigenomic variation between elements of the same family, despite a high degree of sequence similarity. Coupling this information to existing and developing approaches to profile TE transcriptional activity (78) will enable a comprehensive dissection of the epigenetic determinants of TE expression. We also propose that profiling H3K36me3 (which is a robust predictor of transcription (79)) or RNA polymerase II binding using HiChIP and PAtChER is an additional strategy to accurately measure TE transcription. Given the capacity of TEs to regulate gene expression in cis (10), knowledge of which loci are in an active or inactive chromatin configuration will also be invaluable to dissect the landscape of TE-derived gene regulatory elements, and to inform experiments designed to establish causal relationships. By unlocking the access to a large fraction of the genome that has thus far remained hidden, the combination of HiChIP and PAtChER will finally enable a full exploration of the diverse and intriguing epigenetic relationships between TEs and their host genomes.

## DATA AND CODE AVAILABILITY

Hi-MeDIP data have been deposited in NCBI’s Gene Expression Omnibus under accession number GSE182306. LINE-1 bisulphite sequencing data have been deposited in EMBL-EBI’s ArrayExpress under accession number E-MTAB-10895. More details about all the sequencing datasets used in this study can be found in Supplementary Table S1.

PAtChER and all code associated with the manuscript are available in the GitHub repository (https://github.com/MBrancoLab/PAtChER; https://github.com/MBrancoLab/Taylor_2021_PAtChER). Links to processed data tracks, including simulated coverage and mapping accuracy tracks, can be found in Supplementary Table S3.

## ACKNOWLEDGEMENTS

We thank Stefan Schoenfelder for help with setting up Hi-C, and Alex de Mendoza for critical reading of the manuscript; GenoMed, the Genomics Core Facility at IRCAN, for sequencing of the BS-ATLAS-seq libraries; and Stuart Newman for supporting the use of the HPC at University of Essex for the analysis of chromatin loops.

