## Supplementary Figures for "Locus-specific chromatin profiling of evolutionarily young transposable elements"

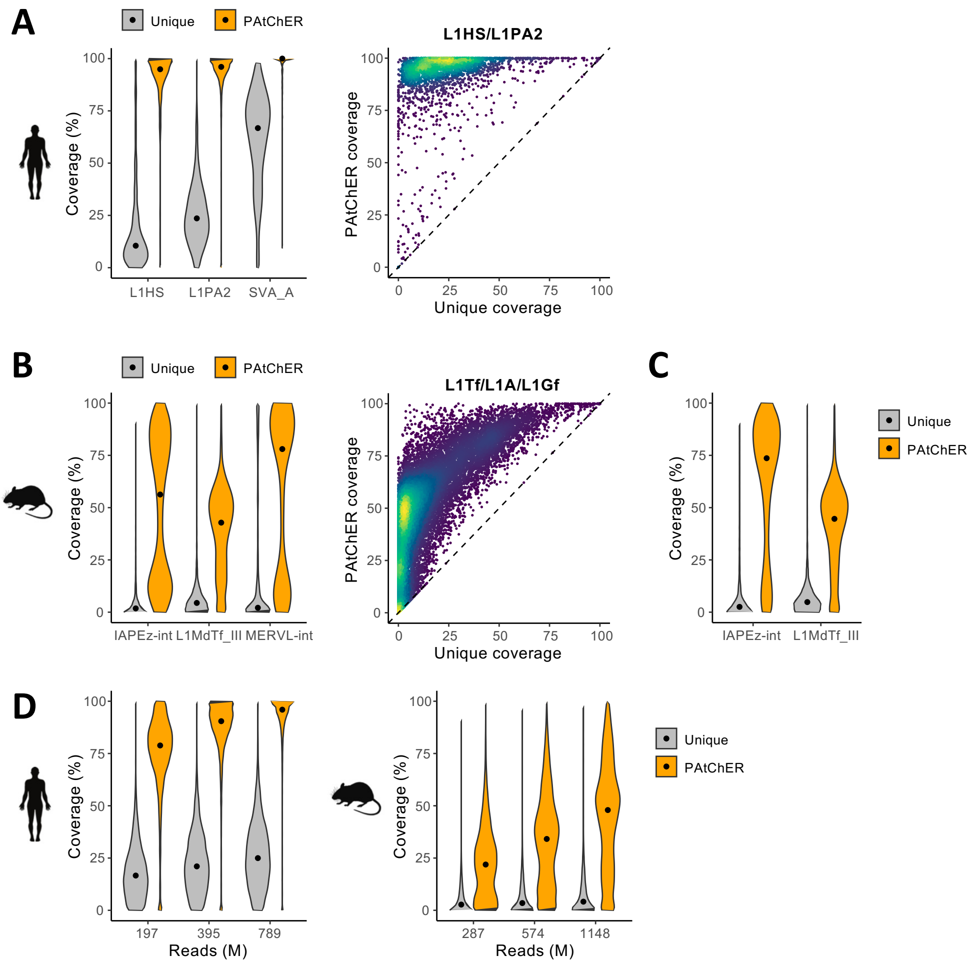


**Supplementary Figure S1** - A,B) Coverage distributions for human (A) or mouse (B) TEs from selected families (elements larger than 1kb). Scatter plots on the right show the coverage of evolutionarily young LINE-1 elements (>1kb). C) Coverage at IAPEz-int and MERVL-int elements after excluding elements known to be absent in the 129 mouse strain (PMID: 22703977). D) Coverage of young repeats (divergence to consensus lower than 3%; length larger than 1kb) as a function of the number of mapped reads.

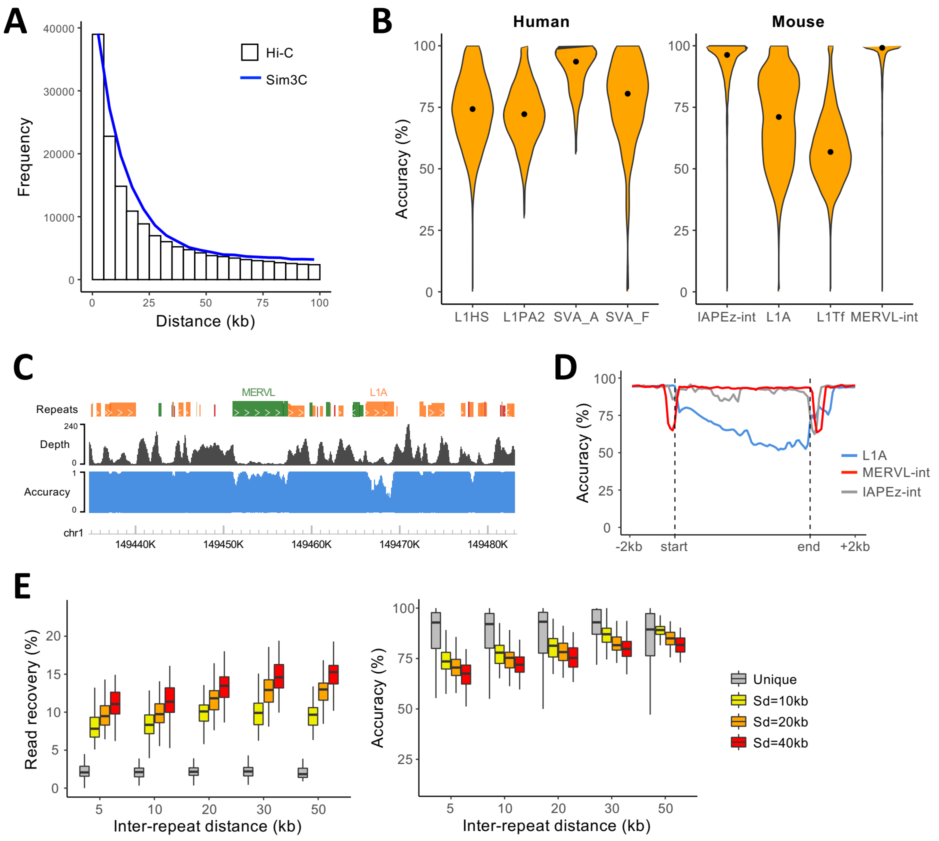


**Supplementary Figure S2** - A) Contact frequency distributions for Hi-C data from mouse ESCs and data simulated by Sim3C (PMID: 29149264). B) Mapping accuracy distributions for different TE families, as assessed from PAtChER-processed Sim3C data. C) Genome browser snapshot of read depth and mapping accuracy tracks for the mouse genome with Sd=10kb, using Sim3C data. D) Average mapping accuracy profile across L1A (>5kb), IAPEz-int (>5kb) and MERVL-int (>4kb) elements, using Sim3C data. E) Hi-C data mapped to the minigenomes were assessed in terms of read recovery (% of reads mapped at original location compared to mm10 alignment) and mapping accuracy (% of reads mapped to original, correct location), as a function of inter-repeat distance, for different values of Sd.

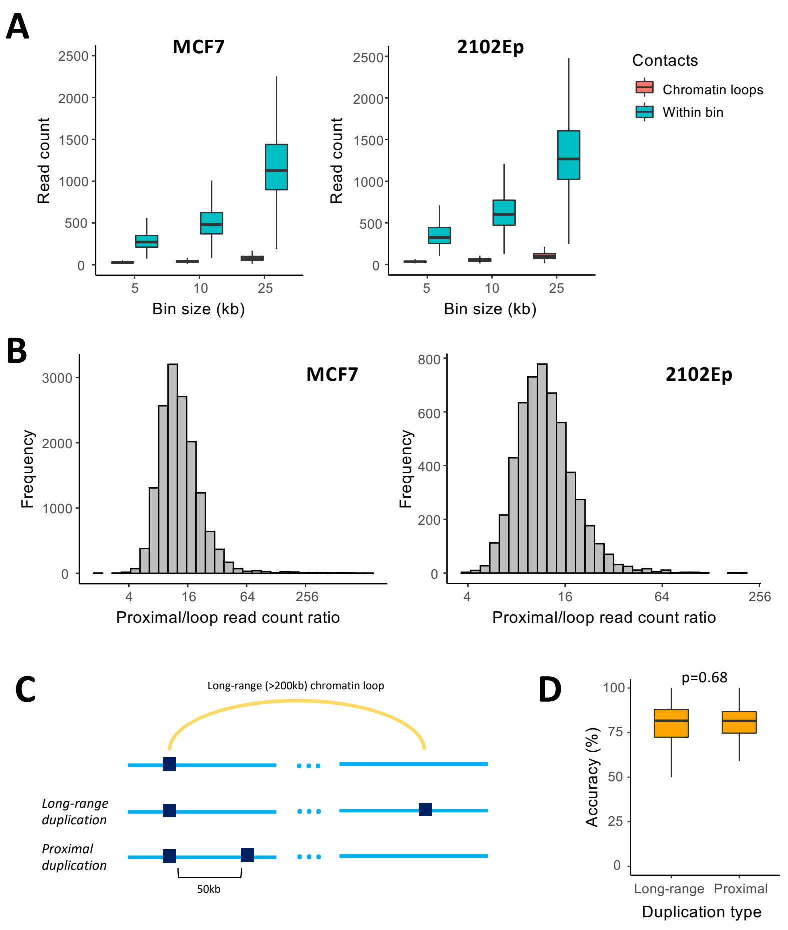


**Supplementary Figure S3** - A) Long-range chromatin loops were detected in Hi-C data from two different cell lines. For each bin at the base of a chromatin loop, the number of reads involved in the loop was compared with the number of reads involved in proximal random contacts within that bin. B) Distribution of the ratio between proximal reads and those involved in a chromatin loop. C) To test the effect of chromatin loops between repeats on mapping accuracy, an artificial minigenome was generated wherein uniquely mappable regions involved in chromatin loops were duplicated at the other end of those loops. This was compared with a minigenome with duplicated regions lying 50kb apart. D) After mapping of MCF7 Hi-C data to the minigenomes, accuracy was measured, with no difference observed between them (Wilcoxon test).

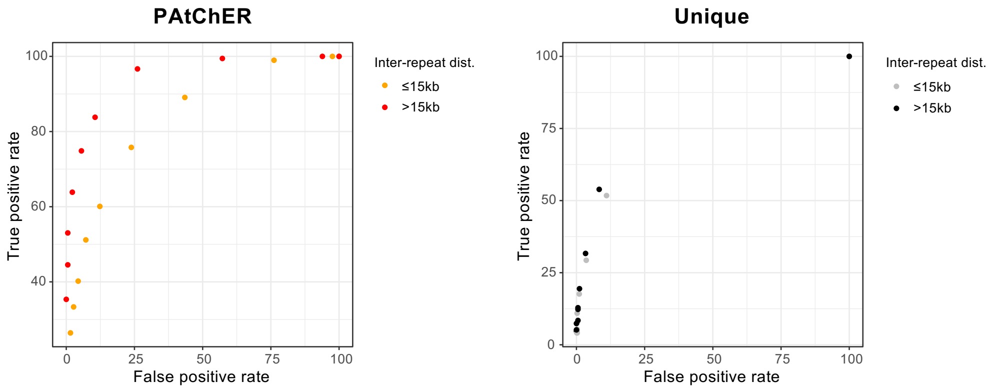


**Supplementary Figure S4** - Receiver operating characteristic curves for H3K27ac peak detection on data mapped to the modified minigenome using PAtChER or uniquely aligned reads. Each data point was generated using a different value for the '-C' parameter of 'macs2 bdgbroadcall'. The true positive rate is the % of mm10 peaks that are found at original location in the modified minigenome. The false positive rate is the % of duplicate locations where peaks were detected.

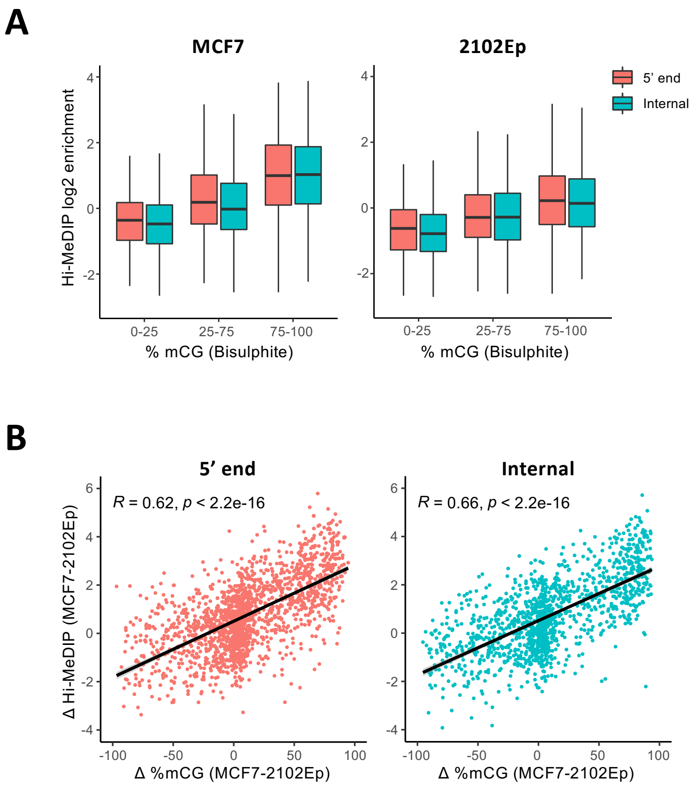


**Supplementary Figure S5** –A) Correlation between PAtChER-processed Hi-MeDIP enrichment values and bisulphite sequencing data (expressed as % of methylated CpGs) in MCF7 and 2102Ep cells, using only all PAtChER-mapped reads. At the 5' end 66% of the reads are unique, whereas in the internal region it is only 29%. B) The DNA methylation difference between MCF7 and 2102Ep cells was calculated using data from either Hi-MeDIP or bisulphite sequencing and compared, using all PAtChER-mapped reads (Pearson's R is displayed).

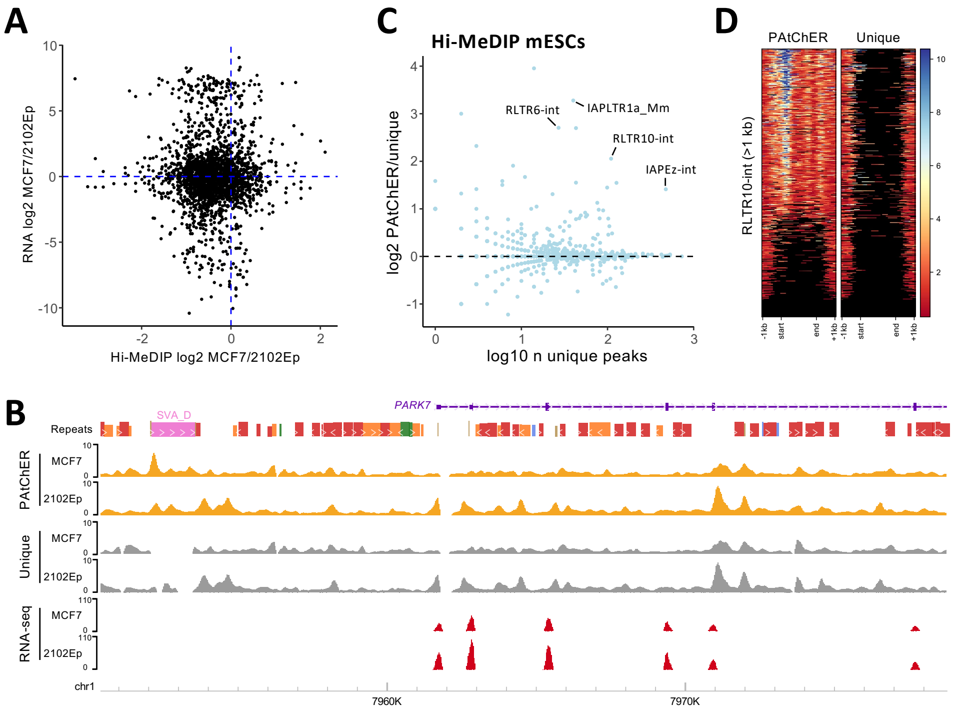


**Supplementary Figure S6** - A) Expression ratio between MCF7 and 2102EP cells for genes within 100kb of an SVA element (larger than 1kb), as a function of the DNA methylation ratio for those elements. B) Genome browser snapshot of the *PARK7* locus, showing a previously characterized SVA element (PMID: 23692647), which PAtChER shows to be differentially methylated between MCF7 and 2102Ep cells, in anti-correlation with the expression of *PARK7*. C) The number of DNA methylation peaks (from Hi-MeDIP) detected in mouse ESCs for each TE family was compared between PAtChER- and uniquely-aligned data. D) DNA methylation enrichment profiles at RLTR10-int elements larger than 1kb in mouse ESCs. Each line in the heatmap represents a single RLTR10-int locus.
